# Four genome assemblies of new wild *Gossypium hirsutum* accessions from the Yucatán Peninsula, the center of domestication

**DOI:** 10.64898/2026.11.02.763792

**Authors:** Corrinne E. Grover, Mark A. Arick, Weixuan Ning, Aya H. Bridgeland, John Z. Yu, Sheron A. Simpson, Monica Wood, Ramey C. Youngblood, Amanda M. Hulse-Kemp, Jonathan F. Wendel, Daniel G. Peterson, Joshua A. Udall

## Abstract

*Gossypium hirsutum* is a leading source of natural cotton fiber, accounting for most of the market share. While domestication and improvement have led to significant gains in fiber quantity and quality, the narrow germplasm base of modern cultivars creates vulnerabilities, including to emerging pests. Wild populations provide a reservoir for genetic diversity that may be valuable in identifying and transferring resistance or stress-adaptation alleles to modern breeding lines. Here we report four new genome assemblies of *G. hirsutum* representing distinct populations from its center of diversity in Yucatán, Mexico. These chromosome-level assemblies are highly contiguous and structurally similar to existing *G. hirsutum* genomes; however, comparative analyses highlight their distinctiveness, suggesting they form a valuable reservoir of untapped diversity. Preliminary analyses of nucleotide-binding leucine-rich repeat (NLR) immune receptor genes underscore this variation, suggesting presence-absence variation may exist among populations. These findings demonstrate the potential of these new genomic resources for discovery of novel disease resistance alleles and, generally, for expanding the narrow germplasm base of the modern cotton cultivars.

## Introduction

Allopolyploid *Gossypium hirsutum* comprises the leading natural textile fiber worldwide, with an estimated demand of ∼119 million bales for 2025-2026 (Meyer and Dew 2026). Agricultural advances in practice and in breeding have increased fiber yield (Fan et al. 2026) such that the present growing year is expected to hit yield records despite a small contraction in global growing areas (Meyer and Dew 2026).

Despite these advances, maintaining or improving yield in crop species with limited genetic diversity faces a number of challenges, including increased temperature and drought risks, emerging pests, and more (Wang et al. 2026). Recent years have seen rising temperatures (Reisinger et al. 2025), which can reduce yield. For example, a small (1°C) maximum temperature increase during specific developmental windows (i.e., flowering) can reduce yield by over 5%, leading to revenue losses (Zahid et al. 2016; Majeed, Rana, et al. 2021; Samuel et al. 2025). Likewise, rising global temperatures have led to higher habitat-associated abiotic stresses, such as low water availability and high soil salinity, which also reduce yield (Lesk et al. 2016; Liu et al. 2025). These stresses are increasing as climate change progresses, resulting in compound effects and leading to greater losses (Sarhadi et al. 2018; Lesk and Anderson 2021). Exacerbating these losses are ongoing and emerging biotic stresses, including tobacco streak virus (Arunachalam et al. 2025), cotton leaf curl virus (Nadeem et al. 2024), and cotton jassid (i.e., two-spot cotton leafhopper; (Cabrera-Asencio et al. 2023)), the last of which has recently spread to North America and may result in yield losses up to 50% in severe cases (Texas Department of Agriculture 2026). Although breeding efforts have targeted many of these stresses, cultivar improvement may have reached a plateau due to both limited and low genetic diversity among accessions (Sakthipriya et al. 2025).

Among solutions for these ongoing and emerging stresses is the potential for incorporating valuable alleles from cotton wild relatives (Hajjar and Hodgkin 2007; Campbell et al. 2010; Kilian et al. 2021; Majeed, Chaudhary, et al. 2021; Tirnaz et al. 2022). Despite historical intentional eradication (Schwendiman et al. 1985), reservoirs of wild *G. hirsutum* remain across the Caribbean, often existing as isolated populations on the leeward sides of many Caribbean islands (Stephens and Phillips 1972; Wendel et al. 1992; Brubaker et al. 1999; Ning et al. 2024; Ning, Hu, et al. 2026), while larger populations occur near the coastline of the northern Yucatán peninsula. These Yucatán populations have recently been confirmed as the center of diversity for *G. hirsutum* (Ning, Arick, et al. 2026), making them invaluable for mining beneficial alleles (Kashyap et al. 2022). Comparative genomics and transcriptomic analyses between crops and their wild relatives have yielded insight into the molecular underpinnings of important phenotypes (Khalid et al. 2025), including variation that may underlie domestication phenotypes in cotton (Wang et al. 2023; Fu et al. 2025). Genomic information for wild *G. hirsutum* is limited to few historical germplasm accessions, either as whole genomes (Li et al. 2024; Yang et al. 2026) or whole genome resequencing data sets (Yuan et al. 2021; Ning, Arick, et al. 2026), the latter having limited value for accurate assessments of gene presence/absence variation and structural variants.

Here we report four new high-quality assemblies for wild *Gossypium hirsutum* accessions collected from the center of domestication, i.e., the Yucatán peninsula. These accessions were sampled from four populations near the northern coastline of Yucatán peninsula, including three sites (Celestún, Sisal, and Centro) on the northwestern Gulf Coast and one (Río Lagartos) on the northeastern coast (Figure 1). Recent resequencing (Ning, Arick, et al. 2026) confirms the wild nature of the populations at these four sites, underscoring their importance in understanding wild *G. hirsutum*, and, by extension, domesticated *G. hirsutum*. We compare these assemblies both among themselves and with existing *G. hirsutum* reference sequences to provide perspective on genomic variability in *G. hirsutum*. We also present an initial analysis of putative nucleotide-binding leucine-rich-repeat receptor (NLR) related genes, noting the potential for differences among accessions derived from closely related populations. Collectively, these demonstrate the utility of these assemblies for understanding diversity among wild *G. hirsutum* and identifying beneficial variation that may be useful in targeted breeding programs.

**Figure 1.**
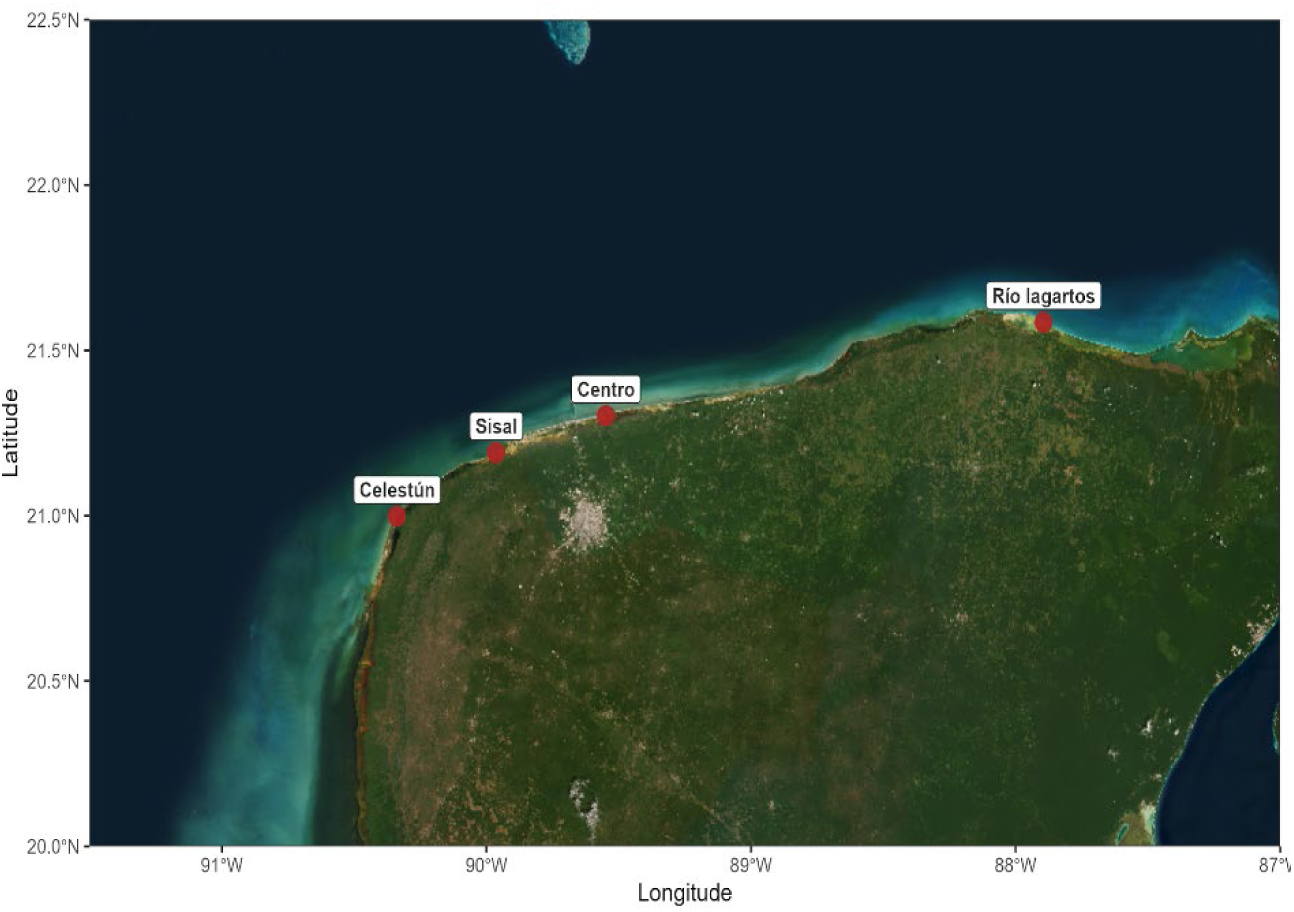
General locations for the four source populations: Celestún, Sisal, Centro, and Río Lagartos. The three northwestern populations represent the center of diversity for wild *G. hirsutum* (Ning, Arick, et al. 2026).

## Methods & Materials

### Sampling, DNA extraction, and genome sequencing

Four wild cotton accessions were selected from a recently published survey of wild cotton diversity (Ning, Arick, et al. 2026) that confirmed the Yucatán peninsula as the center of domestication and diversity for *G. hirsutum*. Field collected seeds from four *G. hirsutum* accessions, i.e., RiChW01 (site: Río Lagartos), CeCoW02 (site: Centro), CePrW01 (site: Celestún), and SiPrW02 (site: Sisal), were germinated on moisturized paper towels in a USDA incubator (College Station, TX USA) kept at 30°C and grown to the cotyledon stage. The dark-growing and sterile conditions in the incubator minimized the amounts of non-nuclear DNA, phenolic compounds, and other biotic contaminants, such as insect residues and microorganisms from greenhouse or field plants.

Fresh young leaf tissue was collected from plants, flash-frozen, and shipped to the USDA-ARS Genomics and Bioinformatics Research Unit (GBRU) in Stoneville, Mississippi for high-molecular-weight (HMW) DNA extraction and PacBio sequencing. HMW DNA was extracted using the PacBio Nanobind PanDNA kit (PacBio, Menlo Park, CA, USA) following the manufacturer’s recommended protocol.

PacBio library construction and sequencing were also performed at GBRU. Genomic DNA was fragmented using a Megaruptor® 3 (Diagenode, Denville, NJ, USA) with a target size of 18–20 kb. SMRTbell libraries were prepared using the PacBio SMRTbell Prep Kit 3.0 according to the standard protocol. Libraries were then size-selected for fragments >10 kb using the PippinHT system (Sage Science, Beverly, MA, USA).

Final libraries were sequenced on the PacBio Revio platform using the Revio Polymerase Kit and SMRT Cell 8M. Raw subreads were processed with the Circular Consensus Sequencing (CCS) algorithm to generate HiFi reads, applying a minimum consensus accuracy threshold of 99.5% to produce high-quality single-molecule consensus sequences.

Hi-C libraries were generated to facilitate chromosome-level assembly of the HiFi reads. DNA from fresh young leaf tissue was used with the Proximo Hi-C kit (Phase Genomics, Seattle, WA, USA) for library generation. Prior to the first step of the kit, leaf tissue was flash-frozen and ground using a mortar and pestle. Library quantification and quality assessment was performed using both a Qubit Fluorometer (Thermo Fisher Scientific, Waltham, MA, USA) and an Agilent 4200 TapeStation system (Agilent Technologies, Santa Clara, CA, USA). Each Hi-C library was sequenced on an Illumina NovaSeq 6000 by Texas A&M Molecular Genomics Core (TAMU-MGC) as paired-end 150 bp reads (PE150) to an average 244M pairs (30.5 x) per library.

### Genome assembly

For each sample, the raw PacBio HiFi reads (average read length 14.7 - 18.0 kb) from each sample (PRJNA1519886; Table S1) were de novo assembled using Hifiasm v.0.19+ (Cheng et al. 2021) to generate a primary contig assembly. The raw assembly was compared to the PSC355 (Cohen et al. 2024) for structure and completeness using a combination of minimap2 (Li 2018) to generate pairwise alignments and dotplotly (Poorten) for visualization. Because each chromosome was represented entirely by <3 contigs, only contigs matching chromosomes from PSC355 were retained for final assembly. For each assembly, scaffolding and comparisons to PSC355 were iteratively performed using their corresponding Hi-C dataset. Hi-C reads were aligned to the assembled contigs using BWA (Li 2013) mem (v0.7.17; parameters -5SP), followed by duplicate marking using SAMBLASTER (Faust and Hall 2014) and subsequent filtering and sorting with SAMtools (Li et al. 2009). Contact pairs were then identified by matlock (Sullivan and Kronenberg) for use in downstream scaffolding. The resulting Hi-C contact links were sorted (LC_ALL=C sort -k2,2 -k6,6) and provided as input to the 3D-DNA pipeline (Dudchenko et al. 2017) to infer contig order and orientation, yielding chromosome-level scaffolds. After scaffolding, the HiC maps were closely inspected for internal inversions and manual curation in Juicebox (Durand et al. 2016).

### Repeat and gene annotation

Each of the four assemblies was aligned to the *G. longicalyx* reference genome (https://www.ncbi.nlm.nih.gov/datasets/genome/GCA_010883175.1/; Grover et al. 2020) using minimap v2.26 (Li 2018) to standardize chromosome number and subgenome orientation. Chromosome orientation was then adjusted using agptools (Ricemeyer et al. 2025) in combination with a Perl script. Genome annotations were performed using two iterations of an EDTA v2.2.2 (Ou et al. 2019) and BRAKER3 v3.0.7.6 (Gabriel et al. 2024) workflow. Briefly, EDTA was initially run without CDS sequences to generate a preliminary genome specific TE library, which was mapped to the genome using RepeatMasker v4.1.2 (Nishimura 2000). The genome was softmasked using bedtools v2.31.1 (Quinlan and Hall 2010) with the mapped repeats; BRAKER3 was then used with the Viridiplantae ODBv11 database (Kuznetsov et al. 2023) to produce a set of preliminary gene annotations. In the second iteration, EDTA was rerun on the genome incorporating the preliminary CDS sequences from BRAKER3 to improve the repeat annotations. Bedtools was again used to softmask the genome before the second BRAKER3 run, which used the updated masked genome and the same (Viridiplantae) protein database.

Completeness for both the genome and annotation of each genome was individually assessed using Benchmarking Universal Single-Copy Ortholog (BUSCO) v5.8.2 (Seppey et al. 2019) in conjunction with the eudicotyledons OBD12 database (Tegenfeldt et al. 2025). LTR Assembly Index (LAI) v3.0.1 (Ou et al. 2018) was also calculated for each assembly using the improved repeat annotations as an additional quality assessment. Gene annotations were filtered based on BRAKER3 transcript scores (retaining transcripts with a score of 0.95 or greater) to minimize low-confidence models while maintaining BUSCO completeness. Final assemblies for each accession are deposited both in NCBI (PRJNA1519886) and CottonGen (https://www.cottongen.org/). The annotations are uploaded to CottonGen. The complete code, including specific parameters, can be found at https://github.com/IGBB/Yucatan-Cotton-Genomes.

### Comparisons among cotton genomes

Assembly quality was assessed using pairwise synteny among the four newly assembled Yucatán genomes (i.e, *G. hirsutum* accessions RiChW01, CeCoW02, CePrW01, SiPrW02), the recently released wild *G. hirsutum* accession TX2094 (Ning, Arick, et al. 2026), and five cultivated *G. hirsutum* telomere-to-telomere assemblies available from CottonGen (Yu et al. 2021): TM-1 (Yan et al. 2025), Jin668 (Xu et al. 2025), YZ1 (Xu et al. 2025), ZM113 (Hu et al. 2025), and TX31 (Fu et al. 2025). Pairwise alignments were conducted through minimap2 and visualized as dotplots using ggplot2 (Wickham 2016) and patchwork (Pedersen 2025), excluding alignments shorter than 100 kb. An additional 157 *G. hirsutum* reference genomes were downloaded from CottonGen (Supplementary Table 1) to estimate their divergence from the four newly constructed reference genomes using PanKmer (Aylward et al. 2023). The additional dataset comprised 125 cultivars, 25 semi-wild accessions, 5 wild accessions, and 2 unknown accessions. All 167 genomes were used in PanKmer to build a kmer index, generate a pairwise adjacency matrix of shared kmers, and visualize the results as a hierarchically clustered heatmap.

PanKmer was also used to estimate the similarity among gene annotations for the four new Yucatán genomes and the previously reported TX2094, the latter being the most comparable of the previously published *G. hirsutum* genomes since it was annotated using the same approach. Orthology among annotations and homoeology of the gene annotations of these five genomes were assessed using GENESPACE v.1.2.3 (Lovell et al. 2022). Because the subgenomes of polyploid *G. hirsutum* are generally independent (Conover et al. 2023), each genome was parsed into the A- and D-chromosomes prior to GENESPACE. Syntenic orthologs among these five genomes (i.e., pangenes) were computed in GENESPACE using ‘query_pangenes’ and riparian plots were visualized using ‘plot_riparian’.

### Survey of putative resistance genes

Putative nucleotide-binding leucine-rich-repeat receptor (NLR) related immune receptor genes were identified and classified using two independent approaches. Protein domain–based classification was performed using Resistify (Smith et al. 2025) on the predicted protein sequences from the genome annotations. A second, complementary approach was used to directly scan the genome sequence for signatures of NLR using NLR-Annotator v2.1b (Steuernagel et al. 2020). Results from each method were integrated, summarized, and visualized in R (R Core Team 2022) using the packages data.table (Extension of “data.frame” [R package data.table version 1.18.4] 2026) and ggplot2 (Wickham 2016). Gene models from each BRAKER3 GTF were intersected with regions identified by NLR-Annotator to identify annotated genes with NLR-Annotator support. The results from the foregoing were combined, and each putative NLR gene was characterized as: (1) identified by both Resistify and NLR-Annotator; (2) identified by Resistify only; or (3) identified by NLR-Annotator only. These results were then intersected with the pangene table produced via GENESPACE to identify orthogroups where an NLR-gene was predicted in at least one genome. Because NLR-Annotator identifies all genomic signatures consistent with NLR genes, category three was restricted to those genes that were identified by NLR-Annotator only but whose orthogroup contained one or more NLR-genes identified by Resistify. NLR clusters in each genome were also identified in R by sorting NLR-genes identified by Resistify (or both) by chromosomal position. Clusters were considered when consecutive NLR genes were identified on the same chromosome and were separated by ≤500 kb.

## Results and Discussion

### Genome assembly and comparison to existing genomes

We report high-quality, chromosome-level genome assemblies for four new wild accessions of the cotton crop species *G. hirsutum* from the Yucatán peninsula (Figure 1): *G. hirsutum* acc RiChW01 from Río Lagartos (site: Chorlitos; hereafter RiCh), *G. hirsutum* acc CeCoW02 from Celestún (site: Consorcio; hereafter CeCo), *G. hirsutum* acc CePrW01 from Centro (site: Pretropical; hereafter CePr), and *G. hirsutum* acc SiPrW02 from Sisal (site: Preseco; hereafter SiPr). Initial assemblies using an average 86.4 Gb (81.6 Gb: Preseco - 98.4 Gb: Pretropical) of PacBio HiFi data for each accession were generally contiguous and similar among accessions. Contig N50 ranged from 86.8 Mb in SiPr to 107.6 Mb in RiCh, with the longest contig ranging narrowly from 127.4 Mb in CeCo to 129.7 Mb in SiPr (Supplementary Table 2). Notably, Hi-C contact maps suggested little scaffolding was required (Supplementary Figure 1) in all four cases, with no scaffolding required for RiCh, consistent with the inbred nature of this population (Ning, Arick, et al. 2026). Total assembly sizes ranged between 2.32 and 2.34 Gb, with uniformly high scaffold N50 values (107.5–107.8 Mb) after scaffolding (Table 1), indicating chromosome-scale contiguity.

**Table 1.**
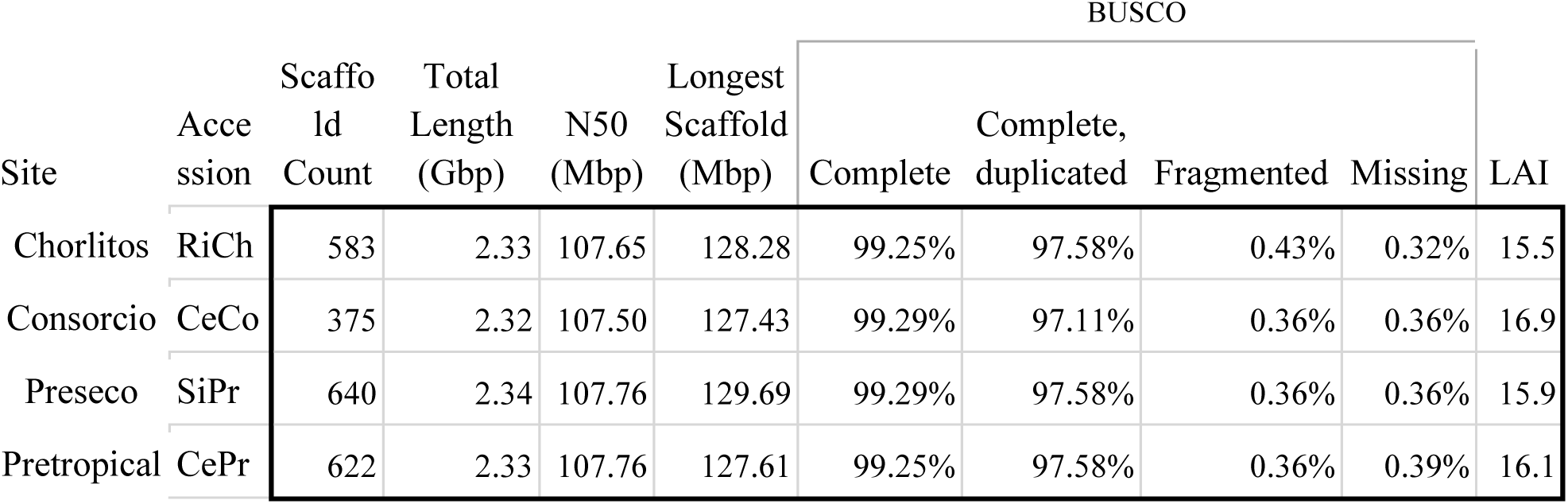
Assembly statistics for four new *G. hirsutum* genomes from Yucatán, Mexico. Scaffold number, total length, and other statistics are given.

|  |  |  |  |  |  | BUSCO |  |  |  |  |
|---|---|---|---|---|---|---|---|---|---|---|
| Site | Accession | Scaffold Count | Total Length (Gbp) | N50 (Mbp) | Longest Scaffold (Mbp) | Complete | Complete, duplicated | Fragmented | Missing | LAI |
| Chorlitos | RiCh | 583 | 2.33 | 107.65 | 128.28 | 99.25% | 97.58% | 0.43% | 0.32% | 15.5 |
| Consorcio | CeCo | 375 | 2.32 | 107.50 | 127.43 | 99.29% | 97.11% | 0.36% | 0.36% | 16.9 |
| Preseco | SiPr | 640 | 2.34 | 107.76 | 129.69 | 99.29% | 97.58% | 0.36% | 0.36% | 15.9 |
| Pretropical | CePr | 622 | 2.33 | 107.76 | 127.61 | 99.25% | 97.58% | 0.36% | 0.39% | 16.1 |

**Supplementary Figure 1.**
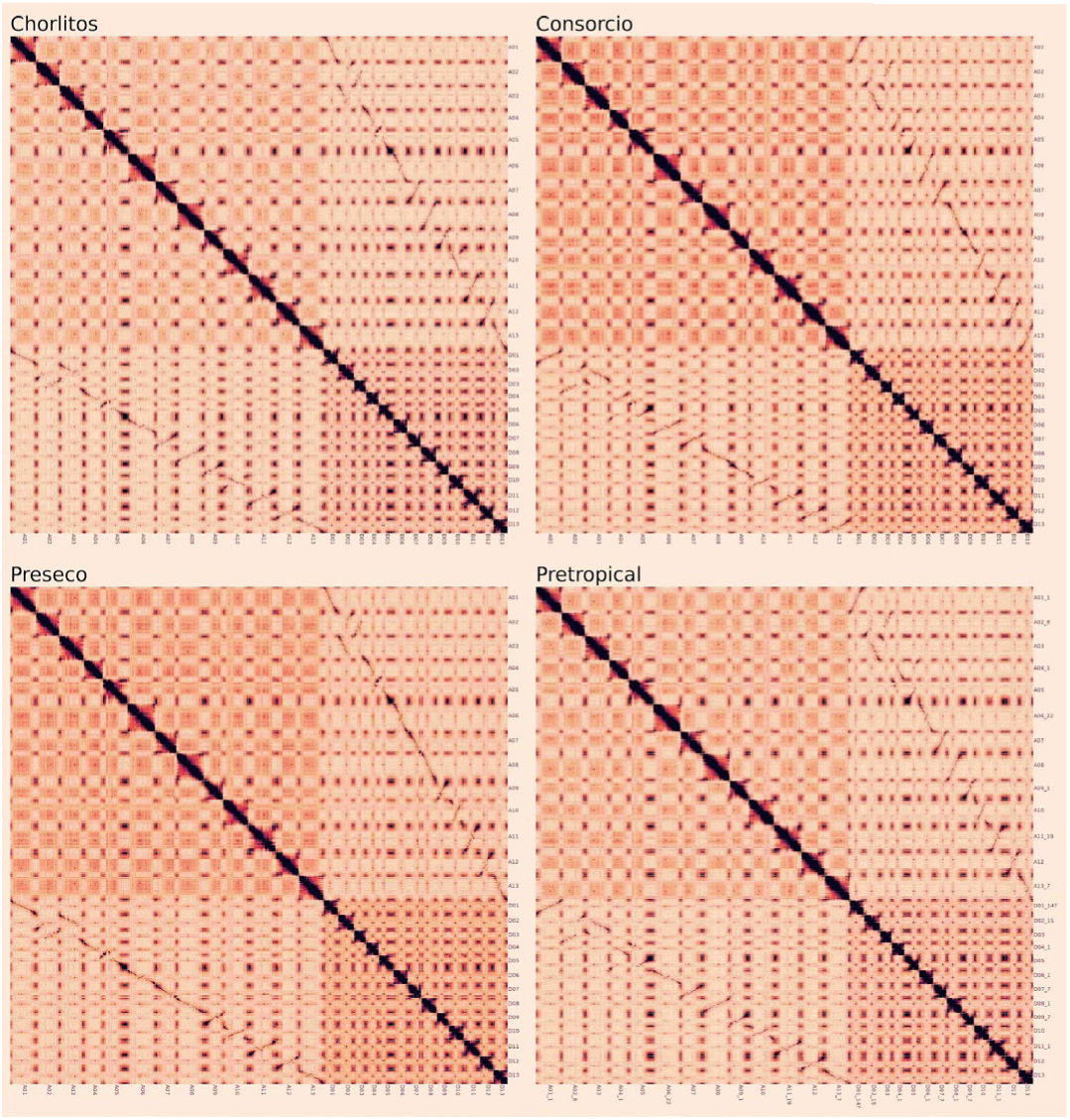
Hi-C contact maps for the final genome assemblies

Although many of the previously published *G. hirsutum* genomes represent modern cultivars, less-domesticated materials have been included in several studies (Li et al. 2024; Fu et al. 2025; Ning, Arick, et al. 2026; Yang et al. 2026). Accordingly, most of the published genomes were expected to be similar with less-domesticated materials exhibiting greater distinctiveness. Here, we used PanKmer to estimate distances among all published *G. hirsutum* genomes along with our newly assembled Yucatán genomes. As expected, most published genomes exhibit exceptionally high kmer similarity, reflecting the narrow germplasm that characterizes domesticated cotton (Ning, Arick, et al. 2026). Among the published genomes, *G. hirsutum* accession ‘Galante 1854’ (HAU_TW100) was also distinct from both the cultivated and wild materials, perhaps reflecting introgression from *G. barbadense* that tends to characterize the Marie-Galante race of *G. hirsutum*. Notably, the four newly sequenced Yucatán *G. hirsutum* genomes (Figure 2), are generally distinct from the remaining *G. hirsutum* genome sequences. The *G. hirsutum* accession TX2094 (Ning, Arick, et al. 2026), also originally sourced from the Yucatán (J. Stewart, pers. comm.), was slightly less distinct from the cultivated materials.

**Figure 2.**
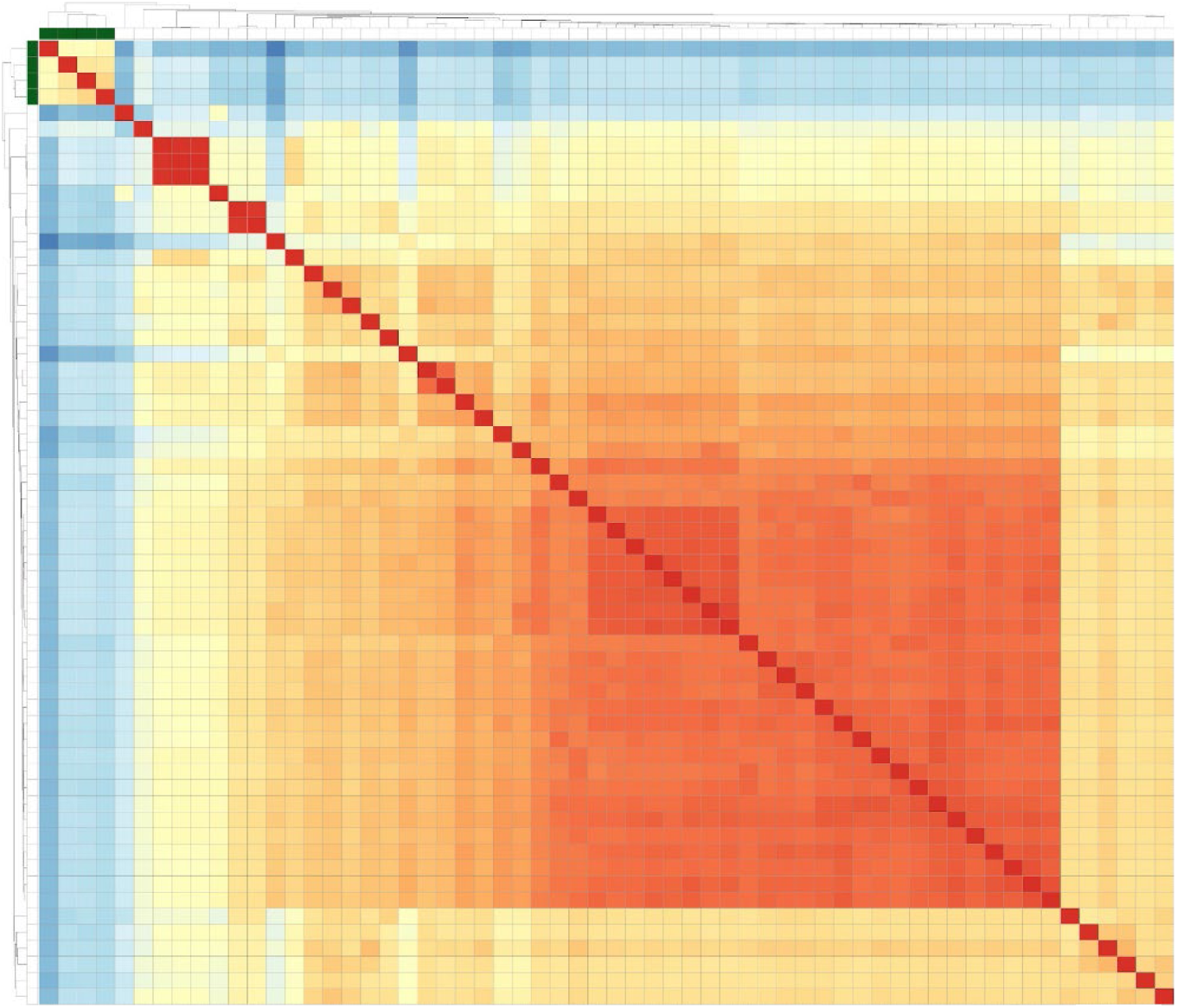
Similarity among a subset of existing *G. hirsutum* genomes and the newly sequenced wild accessions from the Yucatán, denoted by the green bars in the upper left.

We confirmed the quality of each genome using both Benchmarking Universal Single-Copy Ortholog (BUSCO) and LTR Assembly Index (LAI) analysis. BUSCO revealed a high proportion of complete, duplicated BUSCO (97% - 98%; Table 1), as expected for a polyploid species, with an additional ∼2% present as complete, single BUSCO; less than 1% of BUSCO were fragmented or missing (Table 1). LAI likewise confirmed the chromosome-level assembly quality, ranging from 15.5 in RiCh to 16.9 in CeCo. Dotplot comparisons to other recently published *G. hirsutum* genomes, including the telomere-to-telomere releases of *G. hirsutum* cv. ZM113 (Hu et al. 2025) and *G. hirsutum* cv. TM-1 (Yan et al. 2025), reveals high collinearity with existing genomes (Supplementary Figure 2).

**Supplementary Figure 2.**
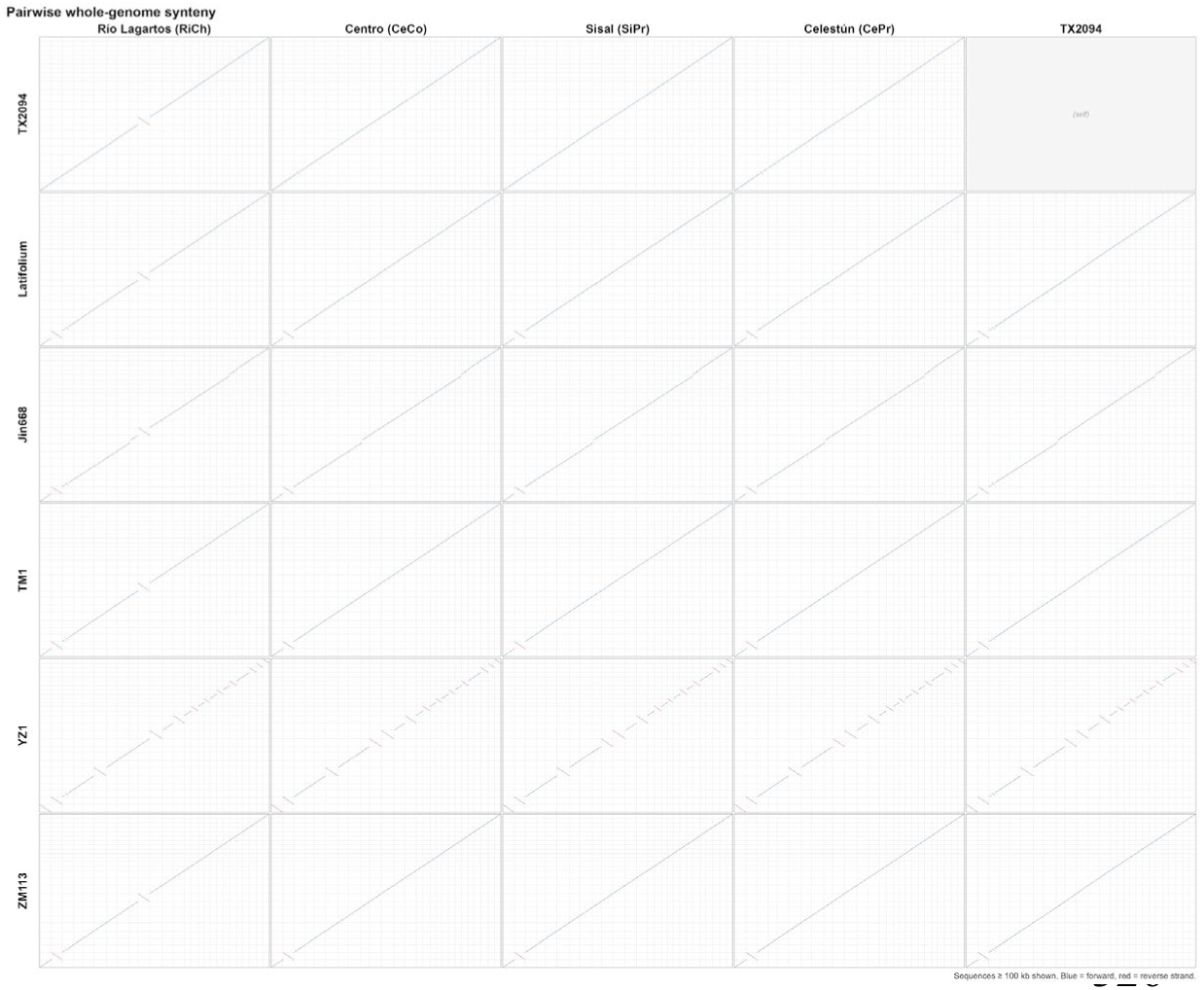
Dotplot comparison between the four newly assembled genomes from the Yucatán (i.e., Chorlitos, Consorcio, Preseco, and Pretropical), the previously reported genome from the Yucatán (TX2094), and five telomere-to-telomere assemblies from CottonGen (Yu et al. 2021): TM-1 (Yan et al. 2025), Jin668 (Xu et al. 2025), YZ1 (Xu et al. 2025), ZM113 (Hu et al. 2025), and TX31 (Fu et al. 2025). Chromosomes are ordered A01-A13 and D01-D13. Only alignments >100kb are shown.

### Genome annotation and orthology

Genomes were *de novo* annotated using a common BRAKER3 pipeline, which predicted between 77,232 to 80,273 total gene models per species (Table 2). Interestingly, three of the four ranged narrowly from 77,232 to 77,620, whereas SiPr had slightly more gene models (80,273) that could neither be attributed to geography nor to genome quality. As expected for a polyploid species, BUSCO analysis of the annotation (Table 2) recovered predominantly complete, duplicated orthologs (94-95%), with around 4-5% occurring as complete, single orthologs; only 0.9-1.3% of BUSCO were either fragmented (0.4-0.6%) or missing (0.5-0.6%). These annotations were approximately evenly divided between the A and D subgenomes of *G. hirsutum* (Table 2), with most genomes, aside from SiPr, having slightly (<1%) fewer genes detected on the A chromosomes. By comparison, the previously reported (and identically generated) annotation of the *G. hirsutum* Yucatán accession TX2094 identified 85,371 predicted coding genes, dispersed as 41,720 genes on the A chromosomes and 43,651 on the D chromosomes (Ning, Arick, et al. 2026). Despite these differences, kmer similarity between the coding sequences among annotations was high (Supplementary Figure 3), suggesting largely consistent annotation among the five genomes.

**Table 2.** Gene and transposable element (TE) annotations for four new cotton genomes. The total number of annotated genes, and the number of annotations per subgenome, is given. BUSCO recovery for the annotation is also provided. Transposable element annotations as listed as copy number (Mbp, % of genome).

|  |  | RiCh | CeCo | SiPr | CePr |
|---|---|---|---|---|---|
| **Gene Annotations** | **Total Genes** | 77,610 | 77,232 | 80,273 | 77,620 |
|  | **A genes** | 38,216 | 38,146 | 40,166 | 38,130 |
|  | **D genes** | 39,394 | 39,086 | 40,107 | 39,490 |
|  | **Complete** | 2,779 (99.1%) | 2,782 (99.2%) | 2,771 (98.8%) | 2,775 (98.9%) |
|  | **Single-copy** | 132 (4.7%) | 144 (5.1%) | 123 (4.4%) | 110 (3.9%) |
|  | **Duplicated** | 2,647 (94.4%) | 2,638 (94.0%) | 2,648 (94.4%) | 2,665 (95.0%) |
|  | **Fragmented** | 13 (0.5%) | 10 (0.4%) | 17 (0.6%) | 13 (0.5%) |
|  | **Missing** | 13 (0.5%) | 13 (0.5%) | 17 (0.6%) | 17 (0.6%) |
|  | **Total searched** | 2,805 | 2,805 | 2,805 | 2,805 |
| **TE annotations** | **LINE** |  |  |  |  |
|  | Jockey | 255 (0.0, <0.1%) | – | – | – |
|  | L1 | 14,733 (11.5, 0.5%) | 16,954 (14.1, 0.6%) | 15,452 (10.3, 0.5%) | 21,904 (13.1, 0.6%) |
|  | **SINE** |  |  |  |  |
|  | tRNA | 56 (0.0, <0.1%) | – | – | – |
|  | **LTR** |  |  |  |  |
|  | Bel-Pao | – | 233 (0.0, <0.1%) | – | – |
|  | Copia | 116,563 (76.5, 3.3%) | 126,607 (80.2, 3.5%) | 135,168 (79.5, 3.5%) | 125,247 (75.3, 3.3%) |
|  | Gypsy | 606,367 (614.5, 26.7%) | 600,364 (555.2, 24.2%) | 585,781 (522.9, 22.7%) | 610,205 (573.5, 25.0%) |
|  | Unknown | 990,420 (674.1, 29.3%) | 1,043,252 (728.0, 31.7%) | 1,047,099 (745.4, 32.4%) | 1,018,877 (720.4, 31.3%) |
|  | **TIR** |  |  |  |  |
|  | CACTA | 103,620 (37.8, 1.6%) | 111,020 (37.7, 1.6%) | 111,918 (38.0, 1.7%) | 78,938 (28.4, 1.2%) |
|  | Mutator | 238,355 (85.0, 3.7%) | 248,226 (79.5, 3.5%) | 241,550 (87.3, 3.8%) | 243,949 (83.9, 3.7%) |
|  | PIF/Harbinger | 149,383 (44.4, 1.9%) | 159,847 (44.1, 1.9%) | 156,821 (43.1, 1.9%) | 165,689 (50.5, 2.2%) |
|  | Tc1/Mariner | 25,180 (11.9, 0.5%) | 26,456 (7.8, 0.3%) | 39,038 (15.1, 0.7%) | 27,304 (7.6, 0.3%) |
|  | hAT | 68,020 (24.8, 1.1%) | 60,458 (22.7, 1.0%) | 68,001 (23.7, 1.0%) | 62,041 (22.4, 1.0%) |
|  | **Non-TIR** |  |  |  |  |
|  | Helitron | 93,573 (21.7, 0.9%) | 102,266 (28.0, 1.2%) | 101,713 (28.8, 1.3%) | 98,918 (27.4, 1.2%) |
|  | **Other** |  |  |  |  |
|  | Low complexity | – | – | – | 140 (0.0, <0.1%) |
|  | Repeat fragment | 157,214 (39.0, 1.7%) | 165,772 (42.2, 1.8%) | 171,908 (44.4, 1.9%) | 160,244 (39.8, 1.7%) |
|  | **Total** | 2,563,739 (1,641.3, 71.4%) | 2,661,455 (1,639.4, 71.4%) | 2,674,449 (1,638.3, 71.3%) | 2,613,456 (1,642.4, 71.5%) |

**Supplementary Figure 3.**
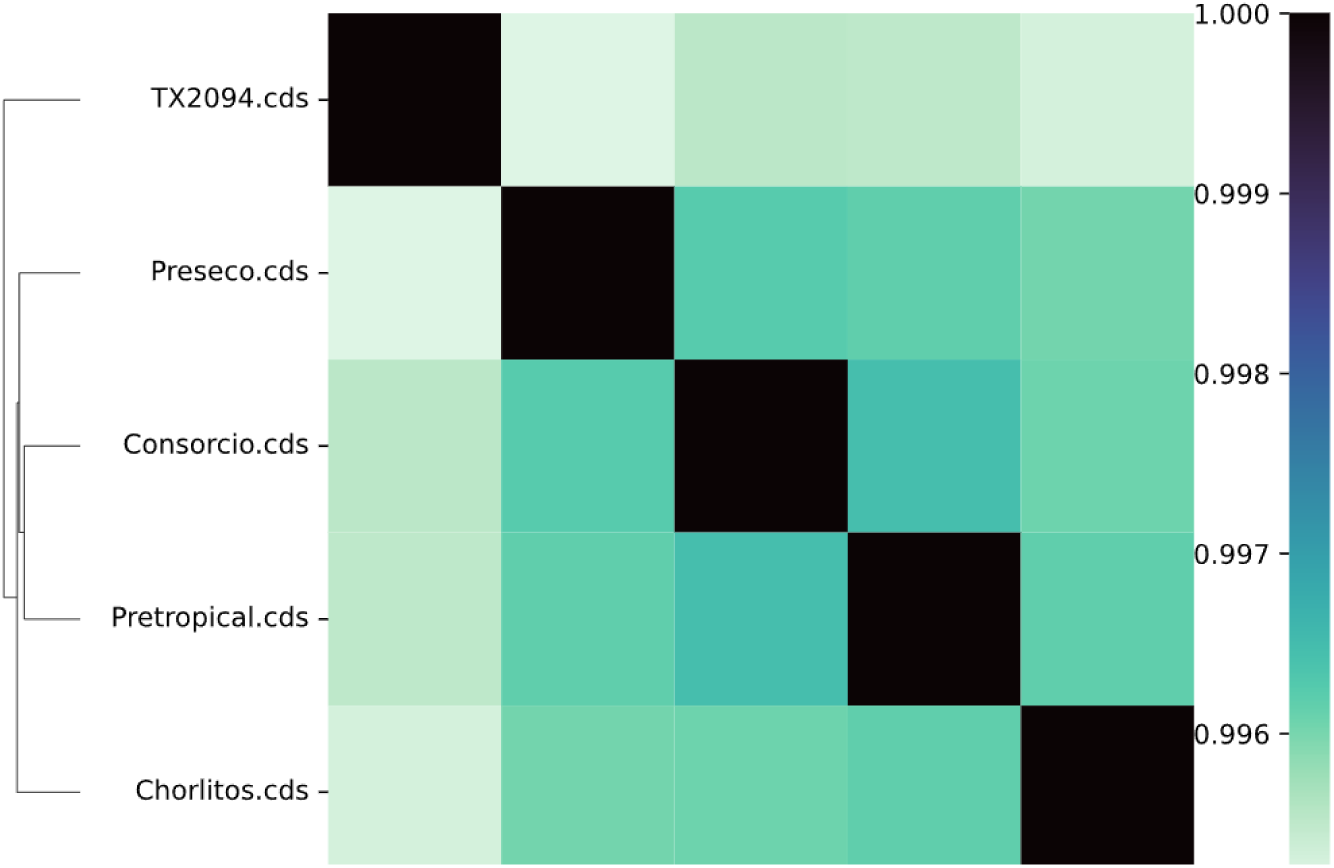
PanKmer similarity among annotations for the four new cotton genomes and the recently published *G. hirsutum* accession TX2094 (Ning, Arick, et al. 2026).

As expected, the genomes of all four assemblies were highly repetitive, with total repeat content ranging narrowly from 71.3-71.5% across accessions with broadly similar repeat profiles (Table 2). LTR-retrotransposons were predominant among repeats, accounting for ∼59% of each genome (∼82% of all annotated repeats). Within this category, unclassified LTR elements were the most abundant, representing 29-32% of each genome, much of which may be composed of degraded *Ty3/Gypsy* LTR-retrotransposons, given their predominance in cotton genomes (Hawkins et al. 2006; Hu et al. 2010; Campos-Dominguez et al. 2025 Mar 21). Identifiable *Ty3/Gypsy* LTR-retrotransposons comprised the second most abundant annotated category (23-27% of the genome), dwarfing the far less abundant *Ty1/Copia* elements (∼3.4%). Class II transposons (i.e., TIR transposons and Helitrons) collectively compose around <10% of each genome, most of whom belong to the Mutator superfamily (3.7% of each genome, on average).

Orthology and homoeology were assessed using these annotations in conjunction with GENESPACE (Figure 3). In general, these genomes were largely colinear, aside from a few large structural changes and known translocations between the two homoeologous subgenomes (Figure 3), e.g., chromosomes 2 and 3, and chromosomes 4 and 5 (Shen et al. 2019). Among the A chromosomes, inversions were found on seven of the thirteen chromosomes (Figure 3). In most cases, these inversions were shared with at least one other accession; however, the NE Yucatán accession RiCh notably had two unshared inversions and one that it shared only with TX2094. The D chromosomes generally had fewer inversions (Figure 3), most of which were shared between accessions. GENESPACE recovered 54,362 orthogroups containing all five accessions, nearly all of which (54,303) comprised single copy orthologs (SCOGs). Orthogroups with genes present in two to four accessions were fewer, although still abundant (two=12,322; three=7,496; four=7,404). Private genes (i.e., those unique to a single accession) were also common (42,743 in total) and approximately evenly distributed, although the number of unique genes in the TX2094 annotation was two-to-threefold greater than the remaining genomes, reflecting its greater number of total annotations. Among the newly generated Yucatán genomes, the number of unique genes ranged from 5442 to 7897, representing 6 - 10% of annotation in each genome.

**Figure 3:**
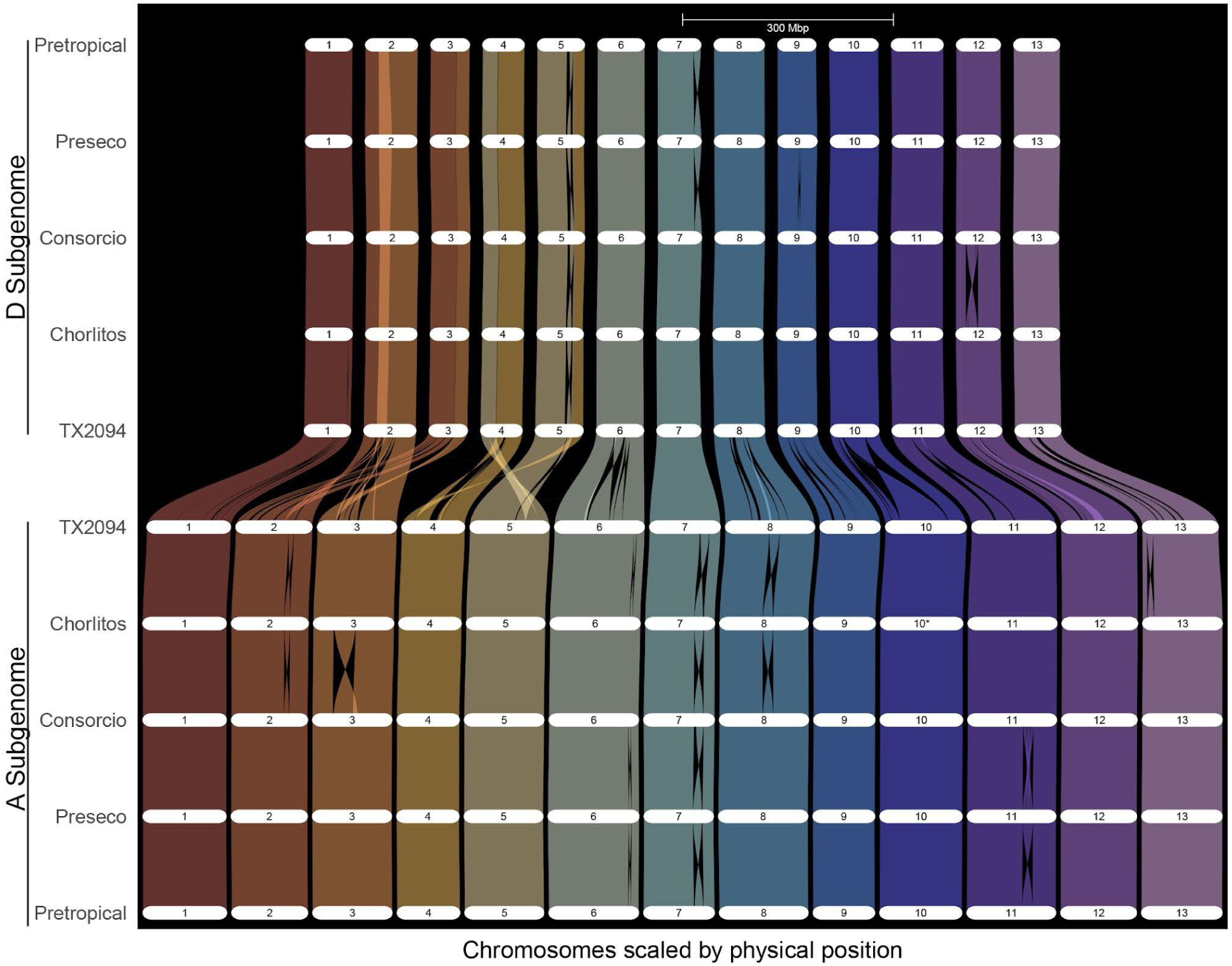
Syntenic relationships among the new Yucatán genomes and the previously published TX2094 genome (Ning, Arick, et al. 2026), which also originated from the Yucatán peninsula and has been maintained by the National Cotton Germplasm Collection (Hinze et al. 2025). Chromosomes from each polyploid subgenome (i.e., At and Dt) were analyzed separately using GENESPACE v1.2.3. Orthologous and homoeologous chromosomal relationships are indicated by colored lines.

### Resistance gene variation among wild accessions

In light of the aforementioned novelty of these new genomes and their genic variability, and given the agronomic importance of stress resistance loci, we explored presence-absence variation (PAV) in putative nucleotide-binding domain leucine-rich repeat (NLR) protein-coding genes, a diverse family of disease resistance genes. Among the five Yucatán accessions analyzed here, we identified 3,103 candidate NLR genes, ranging among accessions from 574 in Pretropical to 652 in TX2094. Among these, nearly 78% of predicted NLRs (2,409) were placed in the GENESPACE pangene matrix (Supplementary Table 3). The majority of these genes were classified into core pangene families (gene n = 1,401; 58%), with NLRs classified into variable (n = 822; 34%) and private (n = 186; 8%) pangene families comprising smaller fractions, indicating modest accession-specific variation. Notably, in most cases, an NLR was predicted in a subset of the genomes containing that pangene (Supplementary Table 3). For example, although the 1,401 individual NLRs assigned to pangene families were classified into nearly as many distinct pangene families (i.e., 1,177 families), nearly half of those pangene families (i.e., 561; Table 3) were considered core. This discrepancy between pangene presence and predicted disease resistance was supported by a combination of both an annotation-based NLR prediction method (Resistify) along with a second, annotation-agnostic (genome-based) analysis (NLR_annotator). Few genes (2; <0.1%) were predicted in the annotation alone and a small proportion (47 loci; <4%) were missed in the annotation but recovered through the genome scan, supporting the observation that only a subset of pangenes display signatures of NLR disease resistance. Rapid turnover of NLR genes has been observed (Michelmore and Meyers 1998; Barragan and Weigel 2021), and we note that orthology is distinct from functional equivalence, which is beyond the scope of the present.

**Table 3.**
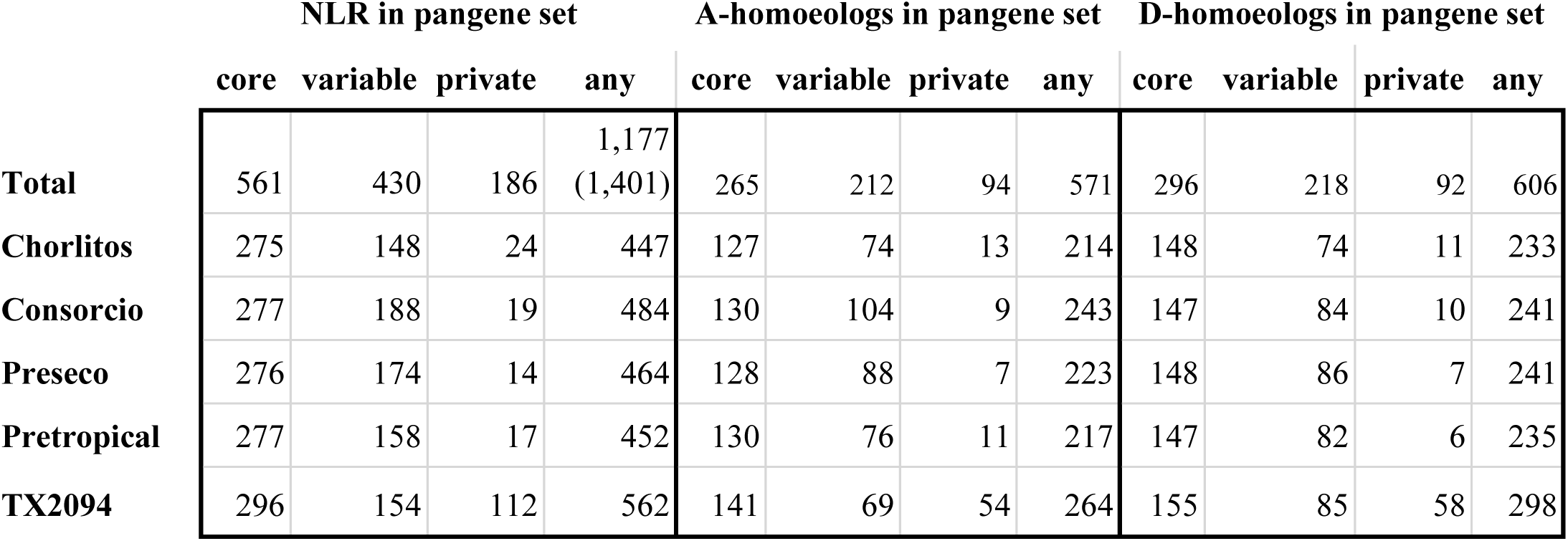
Number of pangene families containing at least one predicted NLR, per category and genome. Core families contain identified orthologs for all five genomes, variable families contain orthologs for 2 - 4 genomes, and private pangenes are found only in one genome. The column "Any" refers to the total number of pangene families in each category that contain an NLR in any genome, and the row "All" refers to the union of NLR gene presence in each genome. Columns 6 to 9 are for those pangenes found on A-subgenome chromosomes only; columns 10 to 13 are for the D-subgenome.

|  | **NLR in pangene set** |  |  |  | **A-homoeologs in pangene set** |  |  |  | **D-homoeologs in pangene set** |  |  |  |
|---|---|---|---|---|---|---|---|---|---|---|---|---|
|  | **core** | **variable** | **private** | **any** | **core** | **variable** | **private** | **any** | **core** | **variable** | **private** | **any** |
| **Total** | 561 | 430 | 186 | 1,177<br>(1,401) | 265 | 212 | 94 | 571 | 296 | 218 | 92 | 606 |
| **Chorlitos** | 275 | 148 | 24 | 447 | 127 | 74 | 13 | 214 | 148 | 74 | 11 | 233 |
| **Consorcio** | 277 | 188 | 19 | 484 | 130 | 104 | 9 | 243 | 147 | 84 | 10 | 241 |
| **Preseco** | 276 | 174 | 14 | 464 | 128 | 88 | 7 | 223 | 148 | 86 | 7 | 241 |
| **Pretropical** | 277 | 158 | 17 | 452 | 130 | 76 | 11 | 217 | 147 | 82 | 6 | 235 |
| **TX2094** | 296 | 154 | 112 | 562 | 141 | 69 | 54 | 264 | 155 | 85 | 58 | 298 |

Nevertheless, the putative PAV among NLR genes from these populations suggests the potential for segregation of novel, and potentially valuable, disease resistance alleles (Barragan and Weigel 2021). Among the accessions analyzed here, the total number of NLR predicted varied modestly from 447 NLR genes predicted in Chorlitos to 562 in TX2094 and more narrowly among the newly sequenced accessions (477 in Chorlitos to 484 in Consorcio). Variation between the subgenomes for each category and genome was generally small (Figure 4), suggesting little to no bias in NLR number or PAV. Although the majority of predicted NLR per genome were assigned to core families (Table 3, and as noted above) each genome typically had 15 to 25 private NLR genes, suggesting the potential for increased gene discovery with additional sequencing. Predicted NLR were unevenly distributed among chromosomes, with chromosomes A07 and D07 having very few predicted NLR whereas Chromosomes A11 and D11 each contained over 150 predicted NLR. As expected, these NLR were generally clustered (Supplementary Figure 4), with several hotspots occurring in the latter half of each chromosome.

**Figure 4.**
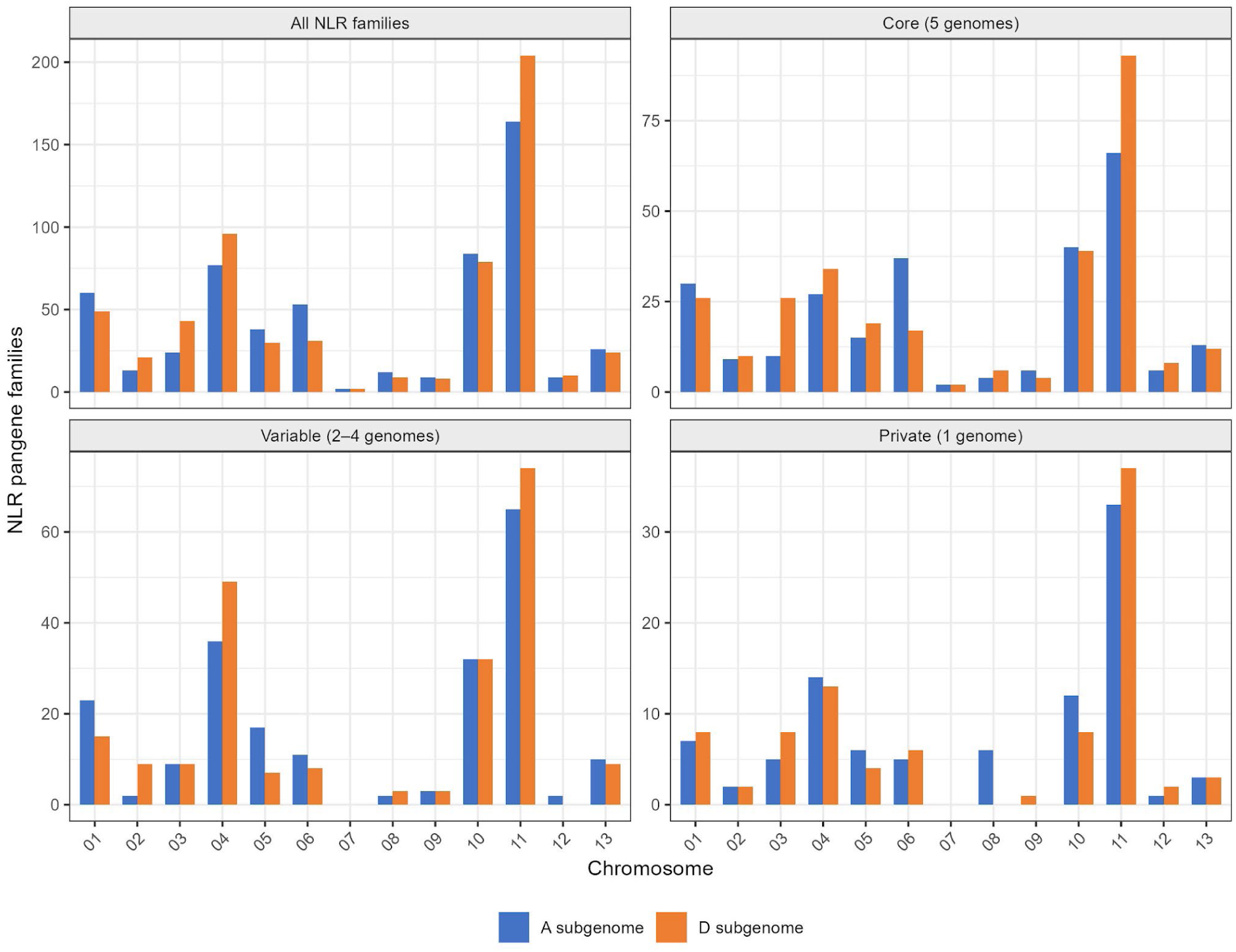
Number of pangene families containing at least one predicted NLR within the pangene set.

**Supplementary Figure 4.**
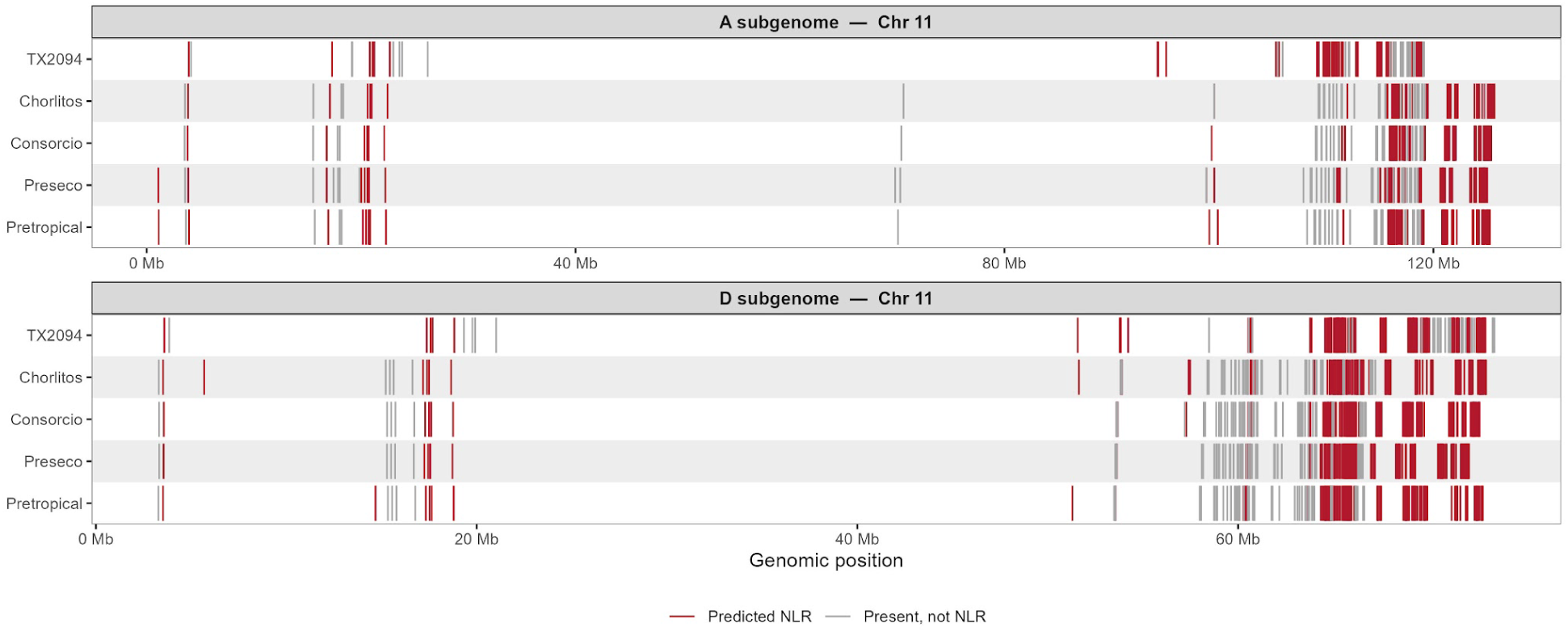
Example of the clustering of NLR genes, as shown on A11 (top) and D11 (bottom). Red lines indicate predicted NLR genes, whereas grey lines are non-NLR orthologs of predicted NLRs.

With respect to NLR class, putative coiled-coil nucleotide-binding leucine-rich repeat (CNL) genes were both the most abundant class and accounted for the largest absolute number of genes assigned to variable or private pangene families. This, however, partly reflects the overall abundance of CNLs; proportionally, putative Toll/interleukin-1 receptor nucleotide-binding leucine-rich repeat (TNL) genes showed a higher fraction of non-core assignments. Among the classes, putative TNL genes were frequently within the variable fraction, potentially indicating greater PAV dynamics in this class. In contrast, putative resistance to powdery mildew 8 (RPW8)-like nucleotide-binding leucine-rich repeat (RNL) genes were rare and almost exclusively assigned to core pangene families, consistent with the conserved role of RNLs as helper NLRs, although, once again, their uniform representation within core pan gene families was also variable. Together, these patterns suggest that CNLs constitute the dominant NLR class in these genomes, whereas TNLs may show greater PAV, providing useful information for future characterization of NLR genes in wild *G. hirsutum* and considerations when screening wild resources for putative valuable disease resistance alleles.

## Conclusion

Here we present four newly assembled genomes representing wild *G. hirsutum* accessions from the center of diversity in the Yucatán. Our preliminary analyses note inversions between the Yucatán cottons and existing *G. hirsutum* genomes, as well as inversions between populations. Furthermore, our results indicate PAV both among these newly assembled genomes and in comparison to existing ones, included PAV among putative NLR-type resistance genes. Importantly, divergence between these newly sequenced genomes and existing genome sequences suggests that these samples may include hitherto unrepresented variation. These genomes provide a foundation for exploring diversity in wild *G. hirsutum*.

## Data availability

The assembled *Gossypium* genome sequences and raw sequencing reads are available at NCBI under PRJNA1519886. Genome sequences and annotations are available from CottonGen (https://www.cottongen.org/) and Zenodo (https://zenodo.org/records/23690962). Relevant code is available at https://github.com/IGBB/Yucatan-Cotton-Genomes.

## Acknowledgements

We thank the Iowa State University ResearchIT unit for computational resources and support. This research used resources provided by the SCINet project and/or the AI Center of Excellence of the USDA Agricultural Research Service, ARS project numbers 0201-88888-003-000D and 0201-88888-002-000D. Claude Code (Anthropic, model Claude Sonnet 5) was used in VSCode (Microsoft) to facilitate the generation of some analytical and data visualization code based upon existing code written by the authors. All analytical decisions and interpretations were made by the authors.

## Funding

This work was funded by the USDA-ARS under 59-6066-4-006 (Genomics of Kenaf, Cotton, and Other Malvaceae to MAA) and 58-3091-5-058 (Genomic Diversity and Genotypic Assessment of Pecan and Cotton to CEG).

## Conflicts of interest

The authors declare no conflict of interest.

